# Growth-inhibiting effects of the unconventional plant APYRASE 7 of *Arabidopsis thaliana* influences the LRX1/FER/RALF growth regulatory module

**DOI:** 10.1101/2023.03.21.533643

**Authors:** Shibu Gupta, Aline Herger, Amandine Guérin, Xiaoyu Hou, Myriam Schaufelberger, Anouck Diet, Stefan Roffler, Thomas Wicker, Christoph Ringli

## Abstract

Plant cell growth involves coordination of numerous processes and signaling cascades among the different cellular compartments to concomitantly enlarge the protoplast and the surrounding cell wall. In *Arabidopsis thaliana*, the cell wall integrity-sensing process involves the extracellular LRX (LRR-Extensin) proteins that bind RALF (Rapid ALkalinization Factor) peptide hormones and, in vegetative tissues, interact with the transmembrane receptor kinase FERONIA (FER). This LRX/RALF/FER signaling module influences cell wall composition and regulates cell growth. The numerous proteins involved in or influenced by this module are beginning to be characterized. In a genetic screen, mutations in *Apyrase 7* (*APY7*) were identified to suppress growth defects observed in *lrx1* and *fer* mutants. *APY7* encodes a Golgi-localized NTP-diphosphohydrolase, but opposed to other apyrases of Arabidopsis, APY7 revealed to be a negative regulator of cell growth. APY7 modulates the growth-inhibiting effect of RALF1, influences the cell wall architecture, and alters the pH of the extracellular matrix, all of which affect cell growth. Together, this study reveals a function of APY7 in cell wall formation and cell growth that is connected to growth processes influenced by the LRX/FER/RALF signaling module.

## INTRODUCTION

The controlled expansion of plant cells requires a plethora of well-coordinated intra- and extracellular processes. These allow for spatially and temporally regulated enlargement of the cell wall surrounding plant cells, a process that requires cell wall material to be synthesized and integrated into the cell wall. Plants are equipped with a refined system to survey and modulate cell wall development by monitoring changes in cell wall composition and architecture. This sensing of cell wall integrity (CWI) involves a number of cell wall proteins and plasma membrane-spanning receptor kinases that help to perceive structural changes and convey this information to the cytoplasm (Voxeur and Hofte, 2016; Vaahtera et al., 2019). Among these, the transmembrane *<u>C</u>atharanthus <u>r</u>oseus* <u>R</u>eceptor-<u>L</u>ike <u>K</u>inase<u>1</u>-<u>L</u>ike proteins (CrRLK1Ls) have been well studied and identified as receptors of growth regulating RALF (Rapid ALkalinization Factor) peptide hormones (Kohorn, 2016; Franck et al., 2018a). RALF peptides induce alkalinization of the apoplast, influence Ca^2+^ dynamics, negatively influence cell growth and modulate plant defence responses (Pearce et al., 2001; Haruta et al., 2008; Stegmann et al., 2017; Abarca et al., 2021). FERONIA (FER) is the best characterized member of the CrRLK1L family of Arabidopsis and is involved in numerous processes including pollen tube perception and rupture in the female gametophyte, plant immune responses, cell growth, and CWI signaling (Escobar-Restrepo et al., 2007; Wolf et al., 2012; Stegmann et al., 2017; Feng et al., 2018; Ortiz-Morea et al., 2022). FER was identified as a receptor of RALF peptides in the context of plant immunity responses and control of cell elongation, where RALF-bound FER causes inactivation of the AHA2 proton pump and, hence, less acidic extracellular pH coinciding with reduced root growth (Haruta et al., 2014; Li et al., 2015). In vegetative tissue, FER functions in a signaling process with Leucine-rich Repeat Extensins (LRXs) to regulate vacuole development, cell growth, salt stress responses, and defence activities (Zhao et al., 2018; Dünser et al., 2019; Herger et al., 2020; Gronnier et al., 2022).

LRXs are extracellular high-affinity binding sites of RALF peptides (Mecchia et al., 2017; Herger et al., 2019; Moussu et al., 2020) tightly linked to the cell wall via their extensin domain that has the typical features of structural HRGPs (hydroxyproline-rich glycoproteins) (Baumberger et al., 2003a). Eleven *LRX* genes are encoded in the Arabidopsis genome, where *LRX8-11* are predominantly expressed in pollen, whereas the others are expressed in the different vegetative tissues. Interaction of most of the LRXs of vegetative tissue with FER has been demonstrated, suggesting an LRX/RALF/FER signaling module (Dünser et al., 2019; Herger et al., 2019; Herger et al., 2020; Gronnier et al., 2022). This is supported by the similar root hair phenotypes of a *fer-4* knock-out mutant and the *lrx1 lrx2* double mutant affected in the root hair-expressed *LRX1* and *LRX2*, respectively (Baumberger et al., 2001; Baumberger et al., 2003b; Duan et al., 2010; Herger et al., 2020). Also, the *fer-4* shoot phenotype is comparable to the shoot-expressed *lrx3 lrx4 lrx5* triple mutant (Draeger et al., 2015; Zhao et al., 2018; Dünser et al., 2019). While the role of LRXs in CWI sensing, cell wall formation, or salt stress is well established, much less is known about proteins that are involved in or modulated by this signaling activity downstream of the LRX/FER/RALF complex.

Cell growth processes are influenced by extracellular ATP (eATP) that is released by plant cells via Golgi-derived vesicles (Cao et al., 2014; Clark et al., 2014). In Arabidopsis, eATP is bound by the plasma membrane-localized L-type lectin receptor kinase LecRK-I.9/DORN1 (DOes not Respond to Nucleotides 1) and the closely related LecRK-I.5/P2K2, subsequently referred to as DORN1 and P2K2, respectively. Mutations in these genes affect eATP-induced Ca^2+^ dynamics and the response to pathogens (Choi et al., 2014; An Quoc et al., 2020). The amount of eATP is not only regulated by the vesicle-mediated export but also by Apyrases (AdenylPYRophosphatASES), nucleotide triphosphate phosphohydrolases that can hydrolyze ATP and other NTPs (Clark et al., 2014). While many organisms, including plants, have ectoapyrases localizing to the cell wall, the seven members of the Arabidopsis Apyrase family localize to the endomembrane system of the Golgi or ER (Parsons et al., 2012; Schiller et al., 2012; Chiu et al., 2015). Downregulation of Golgi-localized *APY1* and *APY2* causes an increase in eATP, indicating that they possibly modify levels of Golgi-localized ATP destined for the extracellular space (Lim et al., 2014). The *APY1/APY2* downregulation also decreases cell expansion and root growth, interferes with pollen tube growth, alters the gene expression profile, and causes modifications of cell wall composition, hence has pleiotropic effects on plant development (Wu et al., 2005; Lim et al., 2014). *apy6 apy7* double mutants have been shown to be affected in pollen grain formation and pollen tube growth including deformation of the exine cell wall (Yang et al., 2013).

Here, we identified APY7 as a modulator of the LRX/FER/RALF signaling module. Mutations in *APY7* alleviate defects in root hair development induced by mutations in *lrx1* and *lrx2* and alter defects developing in *fer* mutants. Opposed to other apyrases, APY7 is a negative regulator of cell growth and appears necessary for the growth-inhibiting effect of RALF1. APY7 influences the sensitivity towards eATP and has an impact on cell wall architecture. These findings reveal functions of APY7, a non-conventional Apyrase with a particular protein structure and atypical effects on cell growth processes compared to other APYs. They also demonstrate that the Golgi-localized APY7 has a significant impact on several aspects of cell growth processes that are connected to the LRX/FER/RALF signaling module.

## RESULTS

### *rol16* suppresses the *lrx1* mutant root hair phenotype

A genetic approach was used to identify genes encoding proteins that play a role in the LRX/RALF/FER signaling module involved in CWI sensing and cell growth. To this end, a suppressor screen was performed on the *lrx1* mutant of *Arabidopsis thaliana* that is impaired in the root hair-expressed *LRX1* gene and develops short, branched, or burst root hairs (Baumberger et al., 2001) (Figure 1A). *lrx1* mutant seeds were treated with the mutagen ethyl methanesulfonate (EMS) and *rol (repressor of lrx1*) mutants were identified in the M2 generation based on a suppressed *lrx1* root hair phenotype. Seedlings of the *lrx1 rol16* mutant identified in this screen show re-establishment of root hair development and, thus, a wild type-like appearance (Figure 1A). Backcrossing of the *lrx1 rol16* mutant with *lrx1* resulted in the F2 generation in a 3 : 1 segregation of *lrx1*: wild type-like root hair formation, indicating that the *rol16* mutation is recessive.

**Figure 1.**
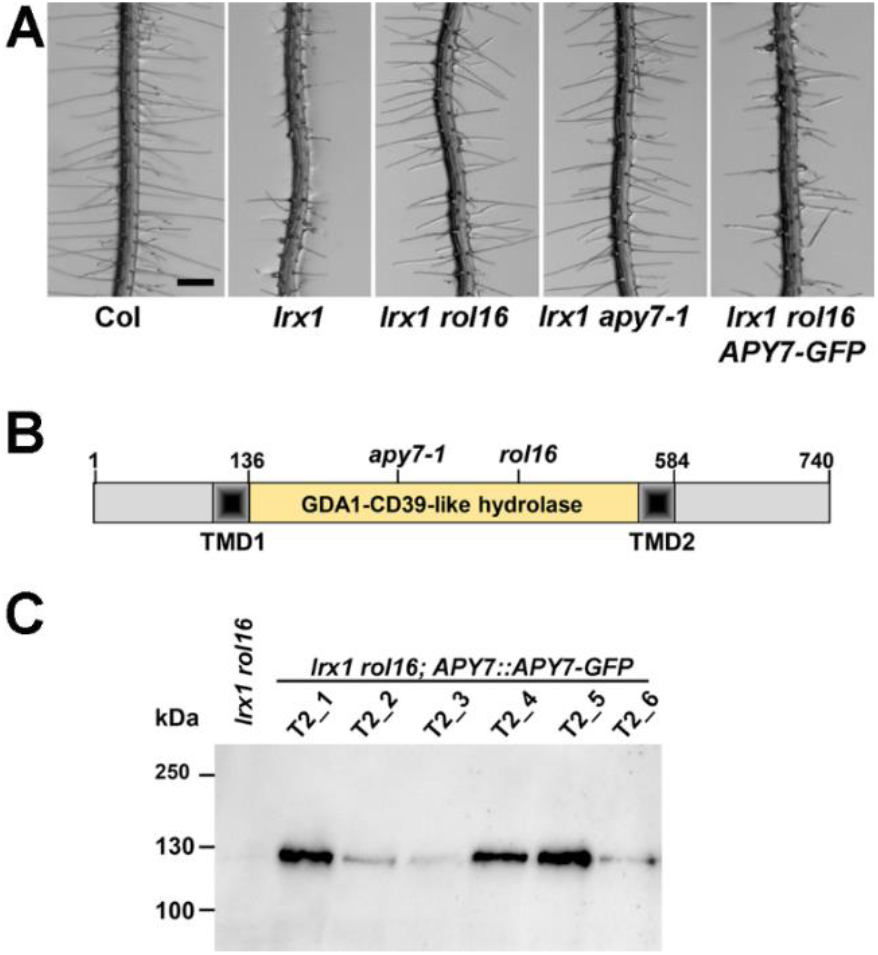
*rol16* suppresses the *lrx1* root hair defect. **(A)** Seedlings grown for six days in a vertical orientation show a defect in root hair development of the *lrx1* mutant compared to the wild type (Col). This defect is suppressed by *rol16* and by *apy7-1*, a second allele of *APY7. APY7::APY7-GFP* successfully complements the *rol16* mutation, resulting in an *lrx1-like* root hair phenotype. Bar=0.5 mm. **(B)** *ROL16* codes for APY7 (Apyrase 7). Apyrases have a nucleoside phosphatase domain, referred to as GDA1-CD39-like domain. APY7 is one of two apyrases of Arabidopsis to have two transmembrane domains (TMD). Positions corresponding to *apy7-1* (SALK_124009) and *rol16* (Trp427stop) on the DNA level are also indicated. **(C)** Total protein extracts of seedlings expressing APY7-GFP were separated by SDS-PAGE and immunoblotted using an anti-GFP antibody. Different T2 lines show different protein levels, all complementing the *lrx1 rol16* mutant phenotype.

### *roI16* is an allele of the ubiquitously expressed *APYRASE7*

To identify the *rol16* mutation, 15 F2 seedlings with wild type-like root hair development were isolated of the segregating population mentioned above, the material was pooled, and DNA was isolated for whole-genome sequencing (WGS). The WGS data of the *lrx1 rol16* mutant was compared to the already existing WGS data of *lrx1* (Schaufelberger et al., 2019) and an SNP with high coverage in the *lrx1 rol16* WGS dataset was identified in the *APYRASE7 (APY7*, At4g19180) gene. A molecular marker for this SNP revealed linkage of the mutation in *APY7* with the *rol16* mutant phenotype in the segregating F2 population mentioned above, suggesting that it causes the *rol16* phenotype. The *rol16* allele contains a G to A mutation introducing a stop codon in APY7 at position Trp427 (Figure 1B). The *apy7-1* allele, containing a T-DNA insertion in *AYP7* at the position corresponding to I263 (SALK_124009) (Yang et al., 2013), was crossed with *lrx1* and the subsequently identified *lrx1 apy7-1* double mutant also suppressed the *lrx1* root hair phenotype (Figure 1A), further supporting the assumption that *ROL16* is *APY7*. Quantitative RT-PCR on total RNA extracted from 10-days-old seedlings revealed a reduction but not depletion of the *apy7* mRNA in the *rol16* mutant compared to the wild type (Suppl. Figure S1A). In view of the interruption of the coding sequence in the mutants, both alleles can be considered loss-of-function alleles despite remaining RNA levels.

APY7 has a GDA1-CD39 (nucleoside phosphatase) domain characteristic of apyrases (Knowles, 2011) that is flanked by transmembrane domains (TMD). In Arabidopsis, APY7 and APY6 both have two TMDs, whereas all other APYs have only one TMD (Figure 1B). Five functionally important sequence motifs conserved among apyrases of diverse origins are also present in APY7 (Suppl. Figure S1B). Furthermore, APY7 has a C-terminal extension of around 110 amino acids not found in any other Arabidopsis apyrase, suggesting that APY7 might have a particular role among the Arabidopsis Apyrase proteins (Clark et al., 2014).

A complementation experiment was performed by transforming the *lrx1 rol16* double mutant with an *APY7::APY7-GFP* construct. Seedlings of the T2 generation of several independent transgenic lines developed *lrx1*-like root hair phenotypes (Figure 1A), demonstrating successful complementation of *rol16*. For detection of the recombinant APY7-GFP fusion protein, one hundred seedlings of each of these T2 families were grown for eight days on half-strength MS plates, pooled, and total protein was extracted. After separation of the proteins by SDS-PAGE, immunoblotting using an anti-GFP antibody revealed varying amounts of APY7-GFP detected among the different T2 lines (Figure 1C), all resulting in a comparable complementation. Together, these results demonstrate that *rol16* is indeed an allele of *APY7*.

The tissue specificity was analyzed in wild-type plants transformed with an *APY7::GUS* reporter construct and in the *APY7::APY7-GFP* transgenic lines. The blue colouring induced by the GUS activity was observed at various stages of plant development and in different tissues including root hairs (Suppl. Figure S2), in line with the previous finding of *APY7* being expressed throughout the plant (Yang et al., 2013).

### *rol16* shows genetic interaction with *lrx1 lrx2* and *fer* mutants

Several LRXs of vegetative tissues, including LRX1, have been shown to function in a signaling process with FER, and the accumulation of mutations in *LRX* genes induces *fer*-like phenotypes (Dünser et al., 2018; Zhao et al., 2018; Herger et al., 2020; Gronnier et al., 2022). As LRX1 and LRX2 function synergistically in root hair development, the *lrx1 lrx2* double mutant shows a severe root hair defect (Baumberger et al., 2003b) comparable to *fer-4* knock-out mutant (Duan et al., 2010). To test whether *rol16* is able to suppress the *lrx1 lrx2* double mutant phenotype, an *lrx1 lrx2 rol16* triple mutant was established. Seedlings of this line indeed develop wild type-like root hairs (Figure 2A). Similarly, *rol16* largely suppresses the root hair defect of *fer-5*, a *fer* allele inducing an intermediate root hair phenotype, (Duan et al., 2010) (Figure 2). In the virtually root hair-less *FER* knock-out allele *fer-4, rol16* allowed more root hairs to successfully enter the elongation phase but they remained shorter than in the wild type (Suppl. Figure S3A). Hence, *rol16* caused a partial suppression of the *fer-4* mutant root hair phenotype.

**Figure 2.**
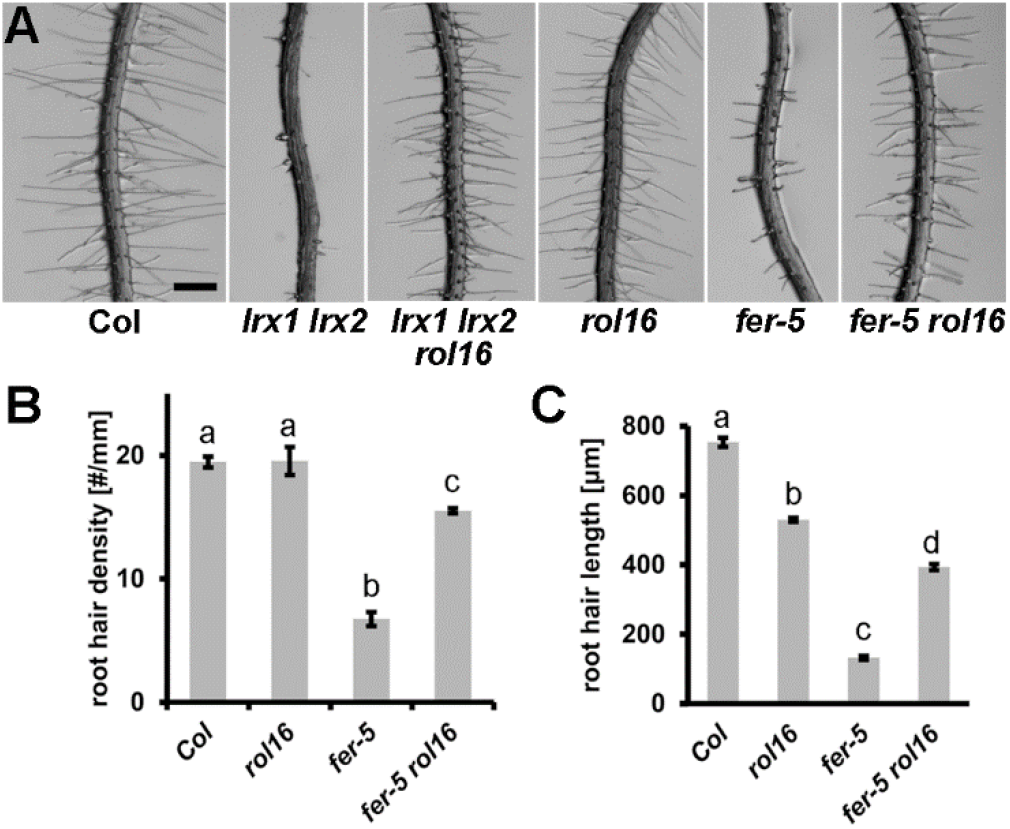
*rol16* suppresses the *lrx1 lrx2* and *fer-5* root hair phenotype. **(A)** Seedlings grown for six days in a vertical orientation show the strong root hair defect in the *lrx1 lrx2* double mutant that is fully suppressed by *rol16. fer-5* contains a T-DNA insertion after the kinase domain and develops an intermediate root hair defect which is suppressed by *rol16*. Bar=0.5 mm. **(B,C)** Quantification of the *fer-5* root hair phenotype and the extend of suppression by *rol16*. Letters above the graphs show statistically significant differences [n>15(B), n>90 (C); student t-test, p<0.001].

In the process of identifying a *fer-4 rol16* double mutant, a distorted segregation of the *fer-4* mutation was observed. Among the progeny of a wild-type plant heterozygous for *fer-4* (*ROL16*^+/+^ *fer-4* ^+/-^), 60 out of 800 seedlings showed a *fer-4* mutant phenotype. This less-than-25% frequency of *fer-4* ^-/-^ was expected considering the function of FER in the fertilization process, which reduces transmission of the *fer* mutation through the female gametophyte (Huck et al., 2003). In a *rol16* mutant population segregating for *fer-4* (*rol16*^-/-^ *fer-4* ^+/-^), however, only 12 out of 800 seedlings were homozygous for *fer-4*. *rol16* does not affect fertilization efficacy, since in the progeny of a heterozygous *rol16* plant, almost one quarter revealed to be homozygous for *rol16* (Suppl. Figure S3B). Hence, the observed strong reduction in the transmission of *fer-4* in the *rol16* mutant background must lay in a synergistic interaction between the two mutations. As a consequence of reduced fertility, *rol16 fer-4* mutants produce shorter siliques with less seeds than the respective single mutants (Suppl. Figure S3C).

In summary, the alleviation of the root hair defects in the *lrx1 lrx2*, *fer-4*, and *fer-5* mutants as well as the impact of *rol16* on fertility of the *fer-4* mutant suggest a genetic relationship between *FER* and *ROL16/APY7* and show that APY7 has an impact on processes that are influenced by LRX and FER proteins.

### APY7 is a negative regulator of cell growth required for RALF1-induced root growth inhibition

The LRX/RALF/FER signaling module modifies cell growth, which prompted us to test whether cell growth is altered in the *apy7* mutants. Indeed, *rol16* and *apy7-1* seedlings grew longer roots than the wild type and had longer epidermal cells (Figure 3A,B). Since LRXs and FER are RALF receptors (Haruta et al., 2014; Mecchia et al., 2017; Zhao et al., 2018; Dünser et al., 2019; Gronnier et al., 2022), it was tested whether APY7 is involved in RALF-mediated growth inhibition. When grown in the presence of 1 μM RALF1 peptide, seedlings of the wild-type Col showed a reduction in root length, whereas seedlings of *fer-4, rol16*, or *apy7-1* mutants showed no inhibition of root growth (Figure 3C). This suggests that APY7 has a function in growth processes that are influenced by RALF1. APY7 negatively influences root growth and cell elongation and is required for RALF1-induced inhibition of cell growth processes.

**Figure 3.**
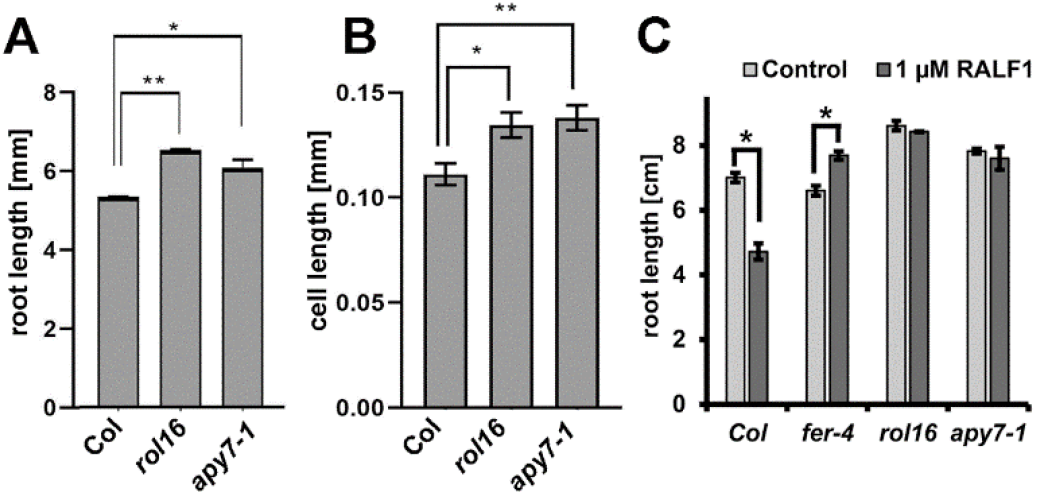
*rol16* and *apy7-1* show increased cell growth and reduced response to RALF1. **(A)** Primary roots length was measured in *rol16* and *apy7-1* mutant seedlings, which were significantly longer compared to the wild type (Col). **(B)** Cell length of trichoblasts was measured in the differentiation zone of the roots and revealed larger cells in the *rol16* and *apy7-1* mutants compared to the wild type (Col). **(C)** Seedlings were germinated on agar plates for four days and subsequently transferred to liquid medium containing 1 uM RALF1 peptide. Wild-type Col seedlings responded to RALF1 by with reduced root growth. *rol16* and *apy7-1* mutants showed insensitivity towards RALF1. Asterisks indicate statistically significant differences in root length (student t-test, n>10, p<0.01). Error bars represent standard deviations (n>5, Student’s t-test, p=* < 0.05, ** < 0.005).

### APY7 influences the apoplastic pH in root tissue

An important effect of RALF1 in regulating root growth is the alkalinization of the apoplastic pH, which inhibits root growth (Haruta et al., 2014; Dünser et al., 2018). Since the *rol16* mutant showed reduced sensitivity towards RALF1, the pH was investigated in *rol16* and *apy7-1* using the ratiometric pH indicator dye HPTS (Barbez et al., 2017). In agreement with previous publications (Barbez et al., 2017; Dünser et al., 2018), an increase in the apoplastic pH was observed in Col seedlings upon RALF1 treatment (Figure 4A,B). Interestingly, even in the absence of RALF1, *rol16* and *apy7-1* mutant seedlings showed a more alkaline pH comparable to Col after RALF1 treatment (Figure 4A,B). Hence, inactivation of APY7 causes a more alkaline pH in the extracellular matrix. This finding urged us to investigate whether there is a correlation between increased pH in the cell wall and suppression of the *lrx1* root hair phenotype. To this end, we used a mutant affected in the proton pump *AHA2* important for acidification of the apoplast (Haruta and Sussman, 2012). As expected, *aha2* seedlings revealed to have a more alkaline extracellular pH (Figure 4A,B). However, *aha2* failed to suppress the *lrx1* root hair defect (Figure 4C), indicating that the *lrx1*-suppressive effect of *rol16* or *apy7-1* is not solely based on the more alkaline apoplastic pH of these mutants.

**Figure 4.**
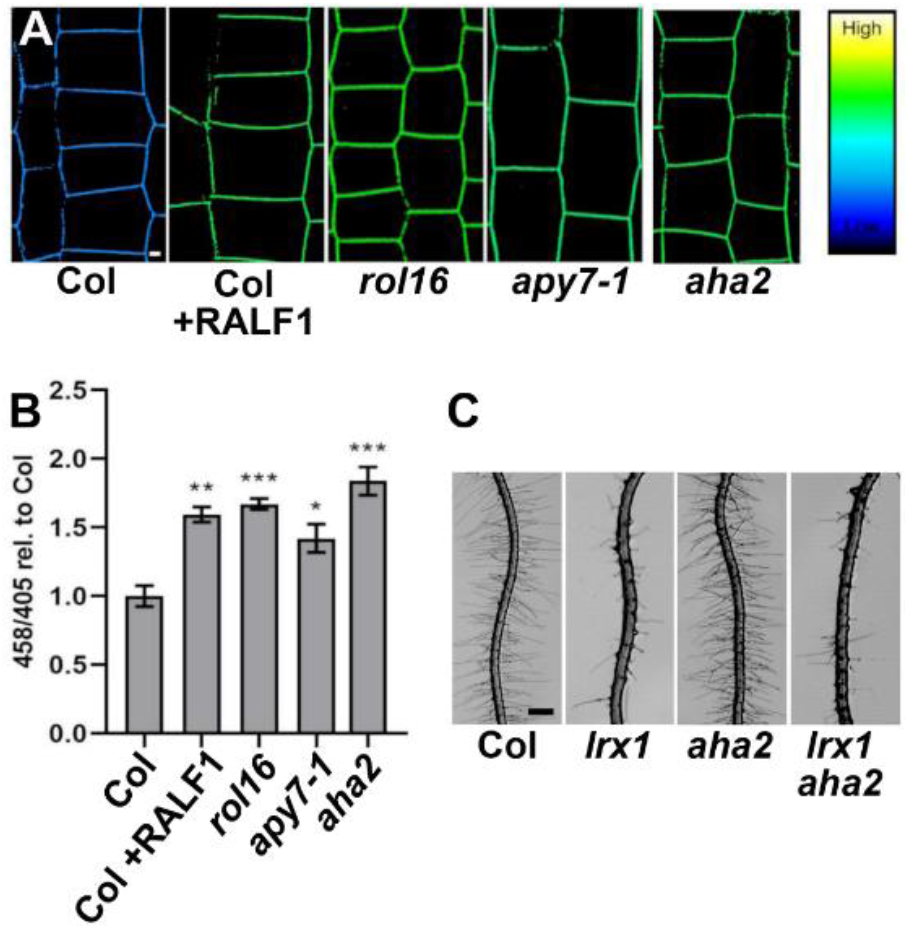
The *rol16* mutants display more alkaline pH in the apoplast. **(A)** Apoplastic pH was measured using HPTS staining in the root epidermal cells in seedlings grown for 4 days on half-strength MS media. Confocal imaging was performed to determine ratiometric pH change. The colour code depicts intensity ratio 458/405 which correlates with pH values. The gradient on the right side represents reference colours for high and low values of pH. Col seedlings were treated with 1 μM RALF1 for 5 minutes and then immediately stained to detect the pH change. **(B)** Quantification of the apoplastic pH in mutants where the y-axis represent 458/405 values of the mutant roots relative to Col that was set to 1. The *rol16* mutant and *apy7-1* display an increase in pH which was significant compared to wild-type Col. The *aha2* mutant also shows more alkaline pH. Ten seedlings per genotype were used as biological replicates. The error bars represent STD. *P-*values were determined by comparing the 458/405 ratios to Col using student t-test: *** < 0.0005, ** < 0.005, * < 0.05. **(C)** The mutation in *aha2* does not suppress the *lrx1* root hair phenotype. Bar = 300 uM.

### Increased sensitivity to exogenous ATP in *rol16* mutant

Apyrases can influence eATP levels as demonstrated for APY1 and APY2 (Wu et al., 2007;

Lim et al., 2014). Levels of eATP produced by Arabidopsis seedlings were measured after growth in liquid medium. Quantification of the eATP levels did not reveal a difference between the *rol16* and *apy7-1* mutants and the wild type (Figure 5A), suggesting that APY7 does not have an obvious impact on eATP levels. To examine whether APY7 is involved in responding to eATP, seedlings were grown on half-strength MS medium containing increasing concentrations of eATP, which induces agravitropic growth/root skewing (Yang et al., 2015) that can be quantified (Suppl. Figure S4). The quantification of agravitropic growth revealed a stronger response of the *rol16* and *apy7-1* mutants to increasing concentrations of eATP compared to the wild type (Figure 5B). Hence, APY7 activity appears to mitigate the response of the root to eATP-induced growth stimulation.

**Figure 5.**
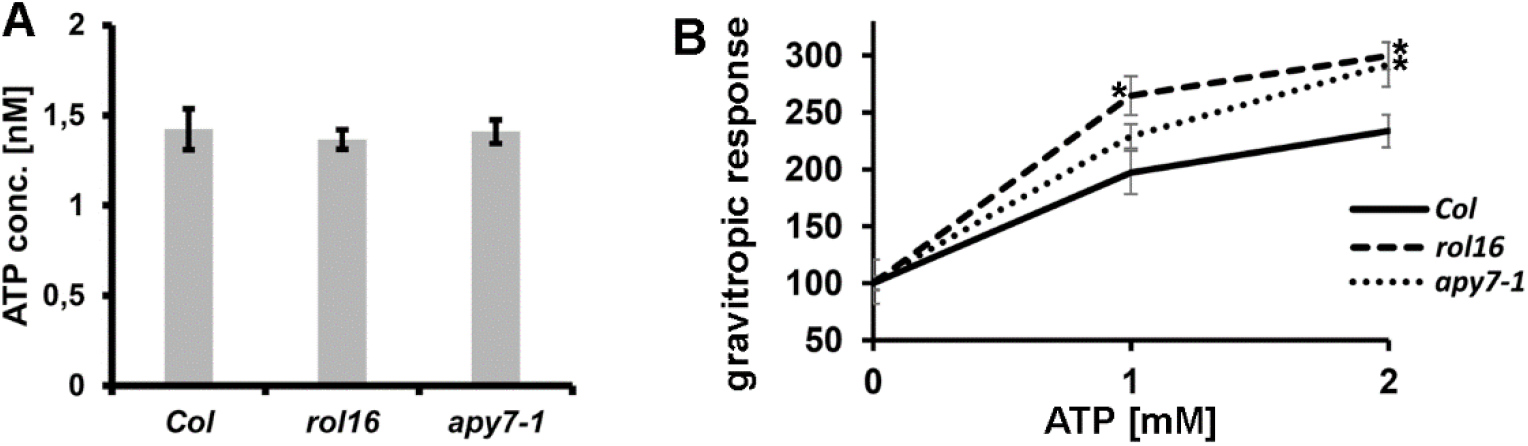
Quantification and response to eATP. **(A)** Extracellular ATP levels in seedlings are not altered in *apy7* mutants. **(B)** Sensitivity to eATP was measured by quantifying the gravitropic response (Grabov et al, 2005) relative to the value in the control sample with 0 mM ATP. The stronger deviation from the gravity vector, the higher the values (see also Suppl. Figure S4). The negative control (0 mM additional eATP) was set to 100% for each line. Asterisks indicate significant differences to the wild type (Col). The *apy7* mutants show a stronger effect upon eATP addition resulting in less gravitropic growth, i.e. stronger agravitropism.

If APY7 is involved in the eATP sensing machinery, mutations in the two plasma membrane-localized ATP receptor proteins *P2K1/DORN1* and *P2K2* (Choi et al., 2014; An Quoc et al., 2020) would possibly also affect the *lrx1* root hair phenotype. An *lrx1 dorn1* double mutant was obtained by crossing the single mutants and showed the same defect in root hair development as the *lrx1* single mutant (Figure 6A). In a next step, the *lrx1 dorn1* double mutant was transformed with a *CRISPR/Cas9* construct targeting *P2K2*. Several lines with insertions of one or deletions of one to several base pair in *P2K2* could be retrieved (for details, see Material and Methods) which all changed the open reading frame, creating a stop codon within 30 amino acids (Suppl. Figure S5). Seedlings of these *lrx1 dorn1 p2k2* triple mutant lines also displayed the typical *lrx1* mutant root hair phenotype (Figure 6A). It was also tested whether DORN1 and P2K2-mediated perception of ATP is required for *rol16*-mediated suppression of *lrx1*. To this end, an *lrx1 rol16 dorn1 p2k2* quadruple mutant was established by crossing of the *lrx1 rol16* and *rol16 dorn1 p2k2* mutants. The *lrx1 rol16 dorn1 p2k2* mutant showed suppression of the *lrx1* mutant phenotype (Figure 6A), indicating that the effect of *rol16* as suppressor of *lrx1* does not depend on functional eATP perception at the plasma membrane. Finally, eATP-induced agravitropism was quantified in Col, *rol16, rol16 dorn1* and *rol16 dorn1 p2k2* lines. Here, the *rol16 dorn1* and *rol16 dorn1 p2k2* lines showed stronger agravitropic growth than *rol16* (Figure 6B), suggesting that the two eATP receptor proteins also influence the gravitropic response. Together, these data suggest that the eATP receptors and APY7 function rather in parallel than in the same linear process to regulate eATP-induced plant development.

**Figure 6.**
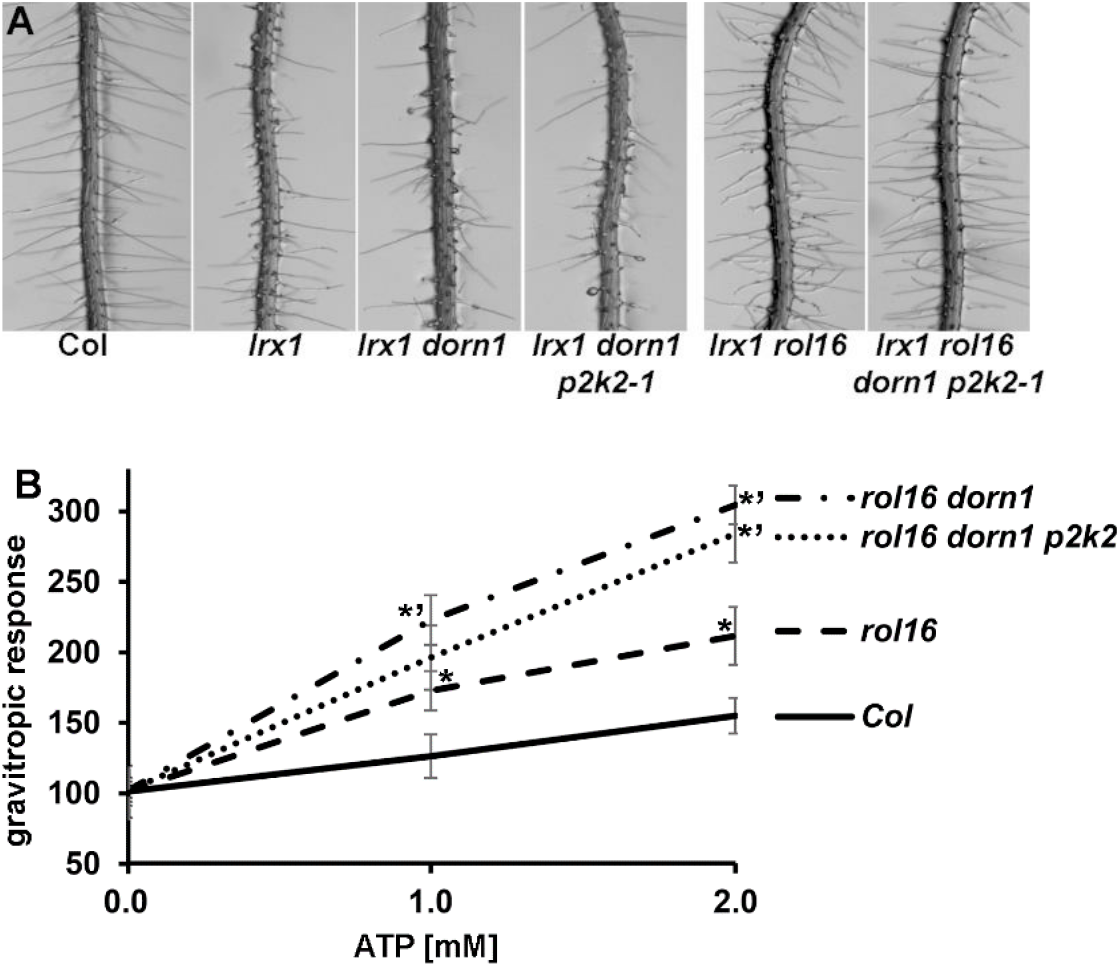
ATP sensing at the plasma membrane does not influence *lrx1* or the suppressive effect of *rol16*. **(A)** Seedlings grown for six days in a vertical orientation are shown. Mutations in the ATP-receptor genes *DORN1* and *P2K2* do not influence the *lrx1* root hair phenotype nor do they influence the ability of *rol16* to suppress *lrx1*. **(B)** *dorn1* and *p2k2* mutations cause an augmented agravitropic response to eATP. The negative control (0 mM additional eATP) was set to 100%. Significant differences (student’s T-test, P<0.05; n > 8, error bar represent STD) between *rol16* and the wild type are indicated with asterisks (*) and between *rol16* and *rol16 dorn1* or *rol16 dorn1 p2k2* with the asterisks (*’).

This raises the question whether APY7 represents an alternative mechanism by which eATP is sensed at the plasma membrane. Arabidopsis proteome data found APY7 only in the Golgi (Parsons et al., 2012) and this localization was confirmed by *APY7-GFP* expression in onion cells (Chiu et al., 2015). Here, a strongly expressing *APY7:APY7-GFP* line was used to investigate a possible co-localization by crossing with a line expressing the plasma membrane marker protein LTI6b-RFP. No evidence of a significant overlap of GFP and RFP fluorescence at the plasma membrane was observed (Suppl. Figure S6). This suggests that APY7 is involved in the intracellular response to eATP and that this response is independent of the signaling processes induced by the eATP receptors DORN1 and P2K2.

### APY7 influences cell wall structures

The biosynthesis of many cell wall-localized polysaccharides takes place in the Golgi (Zhang et al., 2021). Therefore, we investigated whether APY7 has an effect on cell wall structures which might contribute to the observed alterations in root (hair) development. To this end, seedlings were analyzed using a number of monoclonal antibodies (mAbs) against different cell wall epitopes (Suppl. Table S2), and fluorescence intensities of the FITC-labelled secondary antibody were quantified. Consistently in all experiments, differences in labelling wild-type and *rol16* mutant roots were found with LM10 and LM15, two mAbs binding xylan and xyloglucan, respectively (McCartney et al., 2005; Marcus et al., 2008). In the wild type, LM10 stained root hairs, and LM15 both the root and root hairs (Figure 7A,C) consistent with previous reports. LM15 labelling was stronger in the *rol16* mutant than the wild type, which was confirmed by quantification of the fluorescence (Figure 7C-E). A qualitative difference was found for LM10 labelling, which was found only in wild-type but was undetectable in *rol16* mutant root hairs (Figure 7A,B). While the observation of differences in labelling intensities can also be explained by accessibility of the epitopes, these findings demonstrate that a mutation in *APY7* affects cell wall architecture, suggesting a role of the Golgi-localized APY7 in cell wall formation.

**Figure 7.**
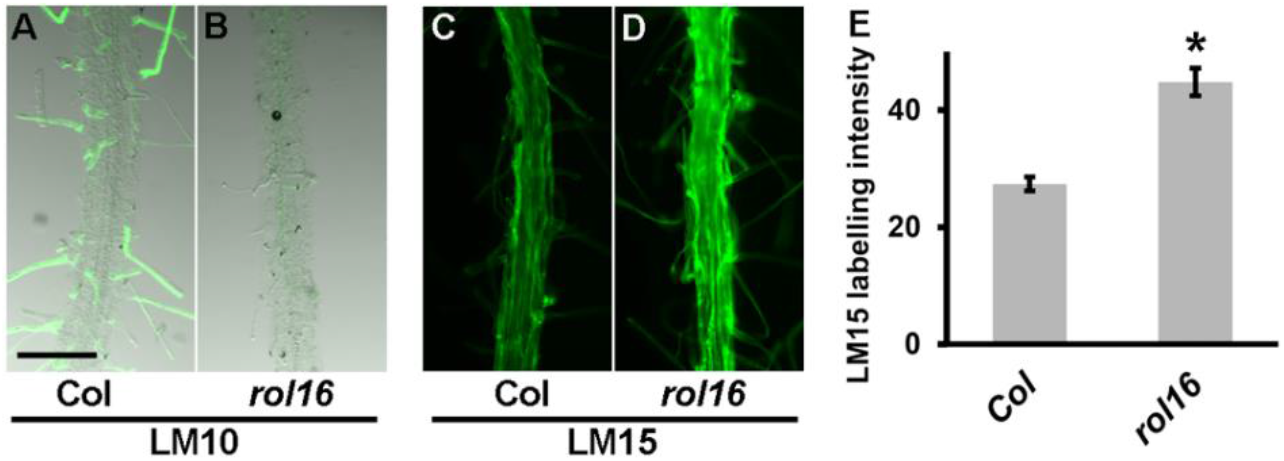
Alterations in cell wall structures in the *rol16* mutant. Six-days-old seedlings were labelled with monoclonal antibodies binding xylan (LM10, **A,B**) and xyloglucan (LM15, **C-E**) epitopes. (**A,B**) Combinations of fluorescence and transmission light microscopy to better visualize root (hair) structures that are only labelled in the wild type. Bar= 0.5 mm. (**C,D**) LM15 labelling along the main root is stronger in *rol16* compared to the wild type, which was quantified by ImageJ (**E**) (student t-test, P<0.01, n>10).

## DISCUSSION

### APY7 is an unconventional apyrase protein

Apyrases degrade NTPs including ATP and thereby can influence many processes including cell growth (Clark et al., 2021). Golgi-localized apyrases regulate levels of ATP required for glycosylations. ATP is also a signaling molecule that is excreted during stress conditions and this extracellular ATP (eATP) is in turn sensed by plants (Cao et al., 2014; Clark et al., 2014). APY7 of Arabidopsis is a particular APY, since its structure with two transmembrane domains is reminiscent of ectoapyrases that localize to the plasma membrane. In Arabidopsis, only APY6 and APY7 have two transmembrane domains and APY7 has an additional and unique C-terminal domain (Clark et al., 2014). The function of this domain and its influence on apyrase activity remains elusive. *APY7* is the only Arabidopsis *APY* not able to complement the yeast apyrase double mutant Δ*ynd1*Δ*gda1* (Chiu et al., 2015), again pointing to a particular activity of APY7. This view is supported by the mutant phenotype of the two *apy7* alleles analyzed here, namely *rol16* and *apy7-1*, that show increased growth. In contrast, mutations in *AYP1* and *APY2* inhibit growth and their overexpression causes increased growth (Steinebrunner et al., 2003; Wu et al., 2007; Yang et al., 2015). While APY1 and APY2 negatively influence eATP levels (Lim et al., 2014), APY7 does not influence eATP levels but rather influences ATP sensitivity. Together, these findings reveal contrasting activities of APY7 compared to other Arabidopsis APYs.

### APY7 is involved in the intracellular response to eATP

In previous studies, application of anti-APY antibodies to pollen tubes increased eATP levels, likely as a consequence of antibody-mediated inhibition of APY activity. This proposes the existence of an APY at the plasma membrane with the enzymatic domain facing the apoplast that reduces eATP levels and is accessible for antibody binding (Wu et al., 2007). APY7 is likely not this plasma membrane-localized APY as it was localized to the Golgi by an APY7-YFP fusion protein and was identified only in the Golgi proteome (Parsons et al., 2012; Chiu et al., 2015). Finally, our analysis did not provide evidence for APY7 localizing to the plasma membrane. It cannot be excluded that small amounts of APY7 evading detection by microscopy are in the plasma membrane. Yet, this contribution would be minor considering that eATP levels appear unaffected in the *apy7* mutants. The increased sensitivity of *apy7* mutants to eATP suggests a role of APY7 in the sensing/response to this signaling molecule. The two lectin-like receptor kinases DORN1/P2K1 and P2K2 are plasma membrane-localized ATP receptors that contribute to the eATP-induced Ca^2+^ response and P2K1 was shown to increase RBOHD-mediated ROS production (Choi et al., 2014; An Quoc et al., 2020). Mutations in these RLKs reduce the sensitivity to eATP, yet neither affect the *lrx1* root hair defect in *lrx1 dorn1 p2k2* triple mutant nor do they prevent suppression of *lrx1* by *rol16* in an *lrx1 rol16 dorn1 p2k2* quadruple mutant. Hence, the effect of the *rol16* mutation appears independent of *DORN1* and *P2K2*. The increased response of the *rol16 dorn1 p2k2* triple mutant to eATP indicates that APY7 and DORN1/P2K2 likely work in parallel signaling pathway. Hence, APY7 is rather responsible for the sensing or responding to intracellular ATP and is not activated via the eATP receptors at the plasma membrane.

### APY7 is part of the machinery downstream of the LRX/FER/RALF signaling module that alters cell wall composition

LRX1 and LRX2, as other LRXs, are high-affinity binding sites for RALF peptides and function in conjunction with the FER receptor kinase (Zhao et al., 2018; Dünser et al., 2019; Herger et al., 2020). Mutations in *APY7* alleviate to different degrees both the *lrx* and *fer* phenotypes, suggesting that APY7 activity is functionally connected to the LRX/FER/RALF signaling module. This view is supported by the synergistic effect between the *fer-4* and *rol16* mutant in respect to fertility. The observed strong reduction in frequency of homozygous *fer-4* mutants by *rol16* can be postulated to be a consequence of a combination of the *fer-4* mutant failing to induce pollen tube rupture at the synergid cells (Huck et al., 2003; Escobar-Restrepo et al., 2007) and a more vigorous growth of mutant pollen tubes that are less prone to burst upon perception at the synergid. The combination of both effects are a possible explanation for the observed further reduction in successful fertilization.

RALF1 is bound by LRX1 and LRX2 (Herger et al., 2020) as well as FER (Haruta et al., 2014) and induces alkalinization of the growth medium and a reduction in cell growth (Pearce et al., 2001). The observed sensitivity of the *apy7* alleles to RALF1 is consistent with APY7 functioning downstream of the LRX/RALF/FER signaling module. Mutations in *APY7* as well as a reduced *RALF1* expression both result in an increased cell elongation (data shown here and (Bergonci et al., 2014)), correlating with the proposed function of RALF1 and APY7 in limiting cell growth. A possible explanation for the observed reduction in sensitivity towards RALF1 might be that the *apy7* mutants show an increased apoplastic pH, which in wild-type seedlings is caused by a RALF1-induced reduction in AHA-type proton exporters (Haruta et al., 2010). This observed alkalinization of the extracellular matrix of *apy7* mutants, however, is not the cause of suppression of *lrx1* as *aha2* mutants also show a more alkaline apoplast (Haruta et al., 2010) but do not modify the *lrx1* mutant phenotype. The more alkaline pH of the apoplast is also unexpected in connection with the increased cell growth in *apy* mutants, since increased cell elongation is thought to correlate with acidification of the apoplast (Rayle and Cleland, 1992) where wall-enlarging expansin activity is increased. Expansins function non-enzymatically, likely by temporarily detaching cellulose-hemicellulose connections (McQueen-Mason et al., 1992). Interestingly, the two antibodies consistently detecting alterations in cell wall architecture of *rol16* compared to the wild type are directed against hemicelluloses, namely Xyloglucan (XG) and Xylan. These changes in cell wall structures might influence the expansion potential of the cell walls. XG is postulated to interconnect cellulose microfibrils by establishing “hotspots” on cellulose-XG interactions (Park and Cosgrove, 2015) and root hairs with reduced xyloglucan content are shorter than those of the wild type (Cavalier et al., 2008). Also, previous work revealed that RALF1 influences the level of *TCH4* expression (Bergonci et al., 2014), a gene encoding a xyloglucan-endo-transglycosylase that promotes cell (wall) elongation (Fry et al., 1992; Eklof and Brumer, 2010). It is conceivable that the increased XG levels observed in the *rol16* mutant might have an opposite effect resulting in the observed increased cell elongation. Xylan is also a hemicellulose found in all cell walls but particularly abundant in secondary cell walls (Pauly et al., 2013; Larson et al., 2014). Opposite to XG, xylan detection is strongly reduced in the *rol16* mutant. Whether the absence of xylan is compensated for by XG or whether the metabolic flow of xylose is simply redirected to XG biosynthesis remains to be determined. There are certainly other factors in the cell wall that influence its expansion. For one, other enzymes modulating hemicellulose network such as XG-hydrolases, but also PMEs (pectin methylesterases) some of which have an alkaline pH optimum. PME activity can be a signaling to trigger pectin turnover and cause cell elongation (Sénéchal et al., 2015; Sénéchal et al., 2017). Cell wall architecture is a key determinant of cell growth, influencing the physical properties of the cell wall and limiting its expansion. Even if the changes in antibody labelling in the *rol16* mutant are due to changes in accessibility rather than the abundance of the epitopes, they demonstrate a modification of the cell wall in the absence of APY7. The biosynthesis of the polymers takes place in the Golgi, where APY7 is predominantly localized, suggesting that the initial biosynthetic steps can influence the final cell wall architecture. It remains to be shown exactly how APY7 influences cell wall composition, whether this is via regulating the import of the large variety of sugar monomers required for polymerization, regulating levels of ATP (and other NTPs) in the Golgi lumen which in turn influences enzyme activity, or by affecting vesicle transport, to name three of many possibilities. The differing phenotypes of plants with altered APY1, APY2, and APY7 levels strongly suggest that apyrases show considerable differences in their activities, resulting in very different output effects on growth processes.

## MATERIALS AND METHODS

### Plant material, growth, EMS mutagenesis and cloning

EMS mutagenesis on *lrx1* loss-of-function mutant has been previously described (Diet et al., 2004). The mutant lines used for mapping *ROL16* have the Columbia (Col) genetic background. Seeds were surface sterilized by a solution of 1% sodium hypochlorite, 0.03% Triton X-100, washed three times with distilled water, plated on half-strength MS (2% sucrose, 0.6% gelrite; unless stated otherwise), and stratified for 2-3 days at 4°C. The plates were then transferred to the growth chamber with a 16h-light/8h-dark cycle at 22°C and grown vertically. Seedlings were transferred to soil for crosses and propagation, and grown in a growth chamber with the 16h-light/8h-dark cycle at 22°C. For genotyping, the molecular marker for *lrx1* has been described (Diet et al., 2004). For *rol16*, the primers SG23 and SG24 (all primers are listed in Suppl. Table S1) produce a PCR fragment with a *Spe*I site in *rol16* but not the wild type. The *apy7-1* mutant was obtained from NASC (SALK_124009) and the mutation was detected with the primers LbB1 and SG76, the wild-type copy with SG75 and SG76. The *dorn1* mutant was obtained by NASC (SALK_042209) and detected with the primers LbB1 and *dorn1_R*, the wild type with *dorn1_F* and *dorn1_R*.

For complementation analysis, a 1.5 kb of *APY7* promoter was amplified with *APY7Prom_F*/*APY7Prom_R*, digested with SpeI and AscI, and cloned into *pGPTV-Kan* (Becker et al., 1992) where the *GUS* gene was cut out with AscI/SacI and replaced by *GFP* amplified from pMDC99 (Curtis and Grossniklaus, 2003) with *GFP_AscI_F* and *GFP_SacI_R*. The *APY7* CDS was amplified with *APY7_F* and *APY7_R*, digested with AscI and into the *APY7::GFP* clone digested with AscI to obtain. For the *CRISPR/Cas9* mutagenesis, the *pKI1.1* plasmid (Tsutsui and Higashiyama, 2017) digested with AarI was ligated with the *P2K2*-specific double-stranded oligo designed based on CHOPCHOP (Labun et al., 2019) using the primers P2K2_KI1_F/ P2K2_KI1_R. For identification of a *rol16* single mutant, the *lrx1 rol16* line was crossed with wild-type Columbia and seedlings of the F2 population were selected by the established markers for the genotype *Lrx1*^+/+^*rol16*^-/-^.

### Plant transformation and *CRISPR/CAS9*-mediated mutagenesis in P2K2

Plant transformation was done by floral dipping using Agrobacterium GV3101. T1 seeds of plant transformed with *APY7::APY7-GFP* in the binary vector *pGPTV-Kan* were selected on kanamycin. Several independent T1 transgenic lines were selected and propagated. T1 seeds of plants transformed with the *P2K2*-specific gRNA in the *CRISPR/Cas9* vector *pKI1.1* were selected based on the RFP fluorescence. Cauline leaves of inflorescences of different T1 plants were used to extract DNA and detect Indels in *P2K2* by PCR amplification of the target region with the primers P2K2_F1 and P2K2_R1, followed by sequencing with P2K2_F1.

### Microscopic analysis

Root hair phenotypes were determined with seedlings grown in a vertical orientation for six days, using an MZ125 stereomicroscope (Leica) and images were obtained with a DFC420 digital camera (Leica). *APY7-GFP* localization was done on a Leica Sp5 confocal microscope (Leica, Wetzlar, Germany) equipped with an Argon laser (488 nm) and Diode-Pumped Solid-State (DPSS) laser (561 nm), hybrid detectors and a 63x (N.A. 1.40) oil immersion objective. Fluorescence emissions were filtered between 500 nm and 550 nm for GFP and 580 nm and 700 nm for RFP. In order to obtain quantitative and comparable data, experiments were performed using strictly identical confocal acquisition parameters (e.g. laser power, gain, zoom factor, resolution and emission wavelengths reception). The parameters were selected for low background and no pixel saturation. Intensity measurements and intensity plots were analyzed using ImageJ.

### GUS staining

Seedlings of several T2 families representing independent transformation events containing the *APY7::GUS* construct were grown and stained for GUS activity. To this end, seedlings were vacuum infiltrated in 0.5 mg/ml X-Gluc, 50 mM Na-phosphate pH 7, 10 mM EDTA, 0.1% Triton X-100 and incubated at 37 oC for several hours. GUS staining was subsequently stopped by exchanging the incubation buffer by 70% EtOH. Picture were taken by a stereomicroscope (Leica).

### RNA extraction and quantitative RT-PCR

Dynabeads^®^ mRNA DIRECT™ Kit 61011 and 61012 (61021) from ThermoFisher was used to extract mRNA directly form grinded leaf tissue. 3μL of bead-bound mRNA was reverse transcribed using the iScript advanced kit (BioRad). qRT-PCR was performed using the primers qRT_apy7_5’_F, qRT_apy7_5’_R, qRT_apy7_3’_F and qRT_apy7_3’_R (Suppl. Table S1). The expression was normalized against the housekeeping genes Act2, EFα and UBQ10 using the following primer pairs: ACT2_F and ACT2_R, EFα_F and EFα_R, UBQ10_F and UBQ10_R (Suppl. Table S1).

### Immunoblotting

Seedlings were grown for 10 days in a vertical orientation on half-strength MS plates, and 100 seedlings were collected and grinded in liquid nitrogen. Proteins were extracted with 200 μl 1% SDS, boiled for 5 minutes, cooled on ice, centrifuged, and 20 μl were mixed with 5 μl 5x Lämmli buffer for loading on a 10% SDS-PAGE gel (BioRad). After blotting was done on nitrocellulose using the semi-dry system (BioRad), the membrane was blocked overnight with 1xTBST, 5% non-fat dry milk powder. Immunolabelling was done in 1xTBST, 1% non-fat dry milk powder with a 1:3’000 dilution of an anti-GFP antibody (Biolegend, 902601), followed by a 1:5’000 dilution of an anti-mouse-HRP antibody (Sigma Aldrich, A 4416).

### Cell wall analysis by immunolabelling

Seedlings were grown in a vertical orientation on the growth medium described above, and incubated in 1x PBS, 3% non-fat milk powder, for 1 h with 10-fold diluted anti-cell wall epitope mAbs and 100-fold diluted secondary, FITC-labelled anti-rat antibody (Sigma F1763). Each antibody incubation was followed by washing three times for 10 min each with 1x PBS. Seedlings were placed on a glass slide containing AF1 anti-fade (Citifluor), covered with a cover glass and fluorescence was analyzed with a thunder microscope (Leica) under non-saturating conditions using a eGFP filter. FITC labelling was quantified by ImageJ and the values for the wild type were set to 1. The microscope and camera settings were maintained for the same mAb, but different mAb required different settings, thus the quantifications of different mAb cannot be compared. Each antibody labelling was done with several seedlings and repeated at least five times. Only consistently observed differences in mAb labelling were considered reliable.

### Growth inhibition assay by RALF treatment

The seedlings were grown for three days in a vertical orientation on half-strength MS plates with 1% sucrose and 1% bactoagar. After three days, they were transferred to half-strength MS (1% sucrose) liquid medium containing 1 μM RALF1 peptide and grown for three more days. The seedlings were laid on agar plate and the root length was measured using Fiji ImageJ software. Same protocol was used for the assays with RALF23 and RALF33. For cell size measurement, the seedlings were grown vertically on half-strength MS plates for 6 days. The epidermal cells in the differentiation zone of primary roots were imaged using Zeiss Axio Imager Z1 microscope. The cell length was measured using Fiji ImageJ software.

### Extracellular ATP analysis

For ATP sensitivity test, the seeds were plated on half-strength MS plates (0.6% gelrite) supplemented with 0 mM, 1 mM ATP, 2 mM ATP and 4 mM ATP. Seedlings were grown vertically for six days, scanned, and the root length and progression on the gravity vector were measured using Fiji ImageJ software to determine the root gravity index (Grabov et al., 2005).

The ATP quantification was performed using ENLITEN^®^ ATP Assay System Bioluminiscence Detection Kit from Promega (protocol by (Yang et al., 2015)). The seedlings were grown in half-strength MS liquid medium for five days and the MS medium was collected. A standard curve was produced with the ATP standard provided in the kit which was used to calculate ATP concentration. Five biological replicates per genotype were used.

### Extracellular pH analysis

The protocol for using HPTS dye has been described in Barbez et al., 2017. The seedlings were grown vertically on half-strength MS media for 4 days. Half-strength MS media containing 1 mM HPTS (Sigma-Aldrich) plates were also prepared for confocal imaging in borosilicate imaging chambers (Lab Tek^®^). The seedlings were stained with 1 mM HPTS dye and immediately used for confocal imaging in the Leica Sp5 microscope. Fluorescence was collected for two forms of HPTS using two different channels: protonated (Excitation at 405 nm and emission peak at 514 nm) and deprotonated (Excitation at 458 nm and emission peak at 514 nm).

A 63x glycerol immersion objective lens was used and the fluorescence data collected was analyzed using Fiji ImageJ software. Five biological replicates per genotype were used for the experiments. Ratiometric image conversion was performed using the Fiji script that was customized by Barbez et al. (2017). Statistical analysis was done using GraphPad Prism8 software.

## ACKNOWLEDGEMENTS

This work was supported by the Swiss National Science Foundation grants Nrs. 31003A_166577 and 310030_192495 (to C.R.) and the University of Zurich.

## Notes

### Competing Interest Statement

The authors have declared no competing interest.

