## Supplementary material for "Growth-inhibiting effects of the unconventional plant APYRASE 7 of *Arabidopsis thaliana* influences the LRX1/FER/RALF growth regulatory module": Suppl. Tables

**SUPPLEMENTARY TABLES**

Suppl. Table S1

| | |
|---|---|
| ***rol16* Primers** | |
| SG23 | CAAGTGCGCTGCACTCTC**A**C |
| SG24 | GCGTTTGATAGGTCTGTTGTTCA |
| ***apy7-1* Primers** | |
| SG75 | CTCCATGTTTTGCGTTTGACC |
| SG76 | CGTGCTTATGATCGGATGGAGA |
| ***APY7::APY7-GFP*** | |
| APYProm_F | TGAACTAGTATTCTAGGACTCTATATGGTCAC |
| APYProm_R | TGAGGCGCGCCTTCTCTTGCCTACATATACTAAC |
| APY7_F | GGCGCGCCATGGTTTTTGGTAGGATCACTG |
| APY7_R | GGCGCGCCTCATTTTGAGCATGTGTG |
| GFP_AscI_F | GGCGCGCCATGAGTAAAGGAGAAGAAG |
| GFP_SacI_R | TCAGAGCTCTTAGTGGTGGTGGTGGTG |
| *dorn1-3* | |
| dorn1_F | CTGAATACTTGCGTCTCCTGC |
| dorn1_R | CAGCTTGCGAGGTTATGATTC |
| P2K2 crispr primers | |
| P2K2_KI1_F | ATTGGCCCGTGTGAAATCCACAGT |
| P2K2_KI1_R | AAACACTGTGGATTTCACACGGGC |
| qRT-PCR primers | |
| qRT_apy7-1_5'_F | TATGCCAATGCGTGTTTCTGAAG |
| qRT_apy7-1_5'_R | TGGAACAGATGATTGTGAACCCG |
| qRT_apy7-1_3'_F | CACTTCCACACCAGCCTAATTTT |
| qRT_apy7-1_3'_R | AAGGGTAAAGAGGAAGAGAGTGA |
| P2K2_qRT_F2 | CCAGCAATGGGACAAGTGGTTCT |
| P2K2_qRT_R2 | GTGAGGCTGCATCGACAACTACA |
| EFα_F | TGAGCACGCTCTTCTTGCTTTCA |
| EFα_R | GGTGGTGGCATCCATCTTGTTACA |

| | |
|---|---|
| ACT2_F | CTTGCACCAAGCAGCATGAA |
| ACT2_R | CCGATCCAGACACTGTACTTCCTT |
| UBQ10_F | GGCCTTGTATAATCCCTGATGAATAAG |
| UBQ10_R | AAAGAGATAACAGGAACGGAAACATAGT |

Primers used in this study. Positions in bold (SG23) indicates SNP compared to genomic DNA to create a CAPS marker with a SpeI in *rol16*.

Suppl. Table S2

| ***mAbs*** | |
|---|---|
| LM 1 | extensins /AGPs |
| LM 2 | AGP |
| LM5 | 1-4 Gal |
| LM 6 | 1-5 Ara |
| LM 7 | partially non-block wise methylesterified HG |
| LM 10 | NRE xylan |
| LM 11 | 1-4 xylosyl |
| LM 15 | xyloglucan |
| LM 18 | partially methylesterified HG |
| LM19 | unesterified HG |
| LM 20 | highly methylesterified HG |
| Jim 7 | partially methylesterified HG |
| JIM 11 | extensin |
| JIM 12 | extensin |
| JIM 13 | AGPs |
| JIM 20 | extensin |

Monoclonal antibodies used in this work. A list of these antibodies, their specificities and list of publications describing them can be found at

https://plantcellwalls.leeds.ac.uk/wp-content/uploads/sites/103/2021/11/JPKab2021.pdf
