## Supplementary material for "Growth-inhibiting effects of the unconventional plant APYRASE 7 of *Arabidopsis thaliana* influences the LRX1/FER/RALF growth regulatory module": Suppl. Figures

Suppl. Figure S1

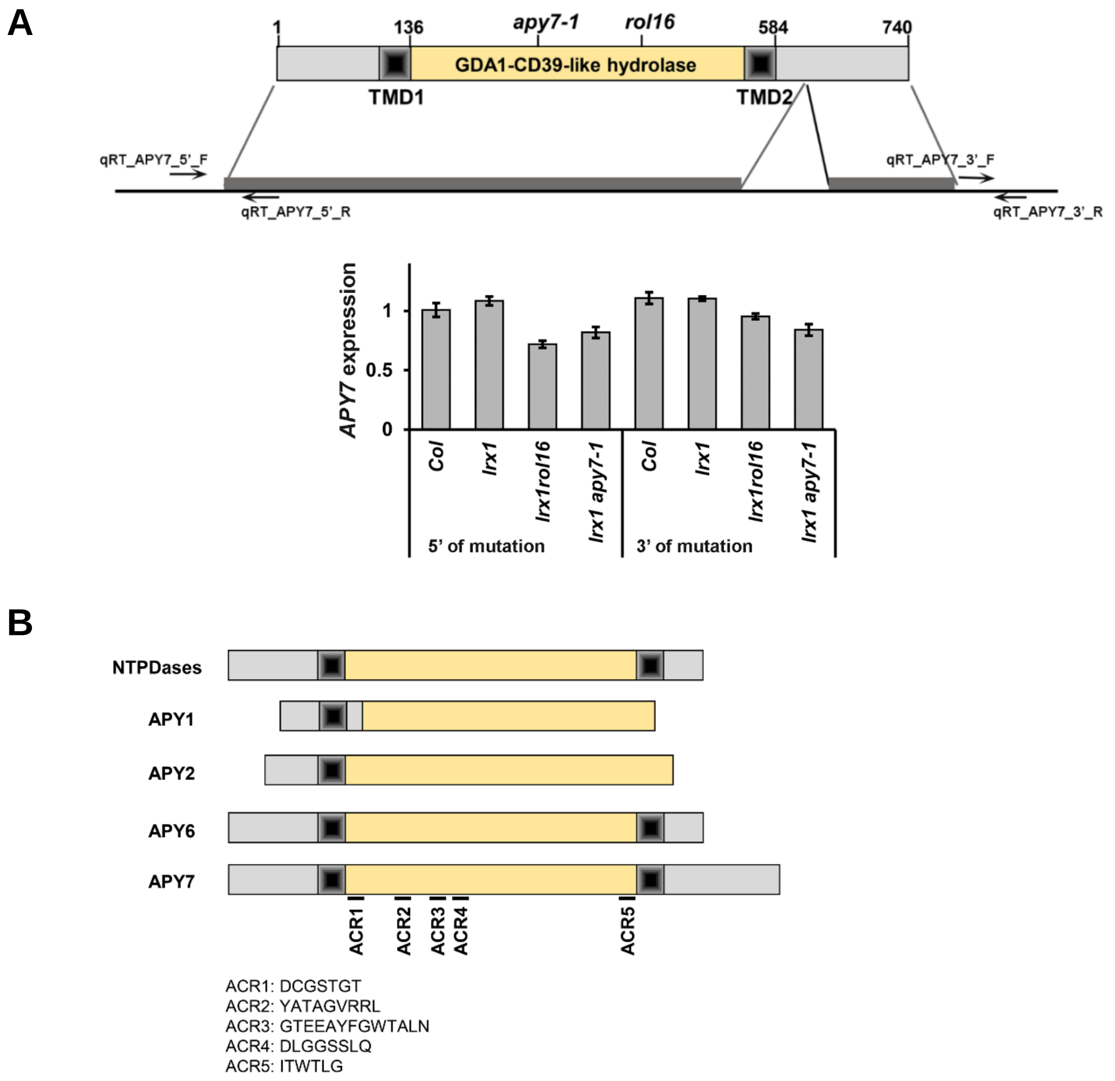


**Suppl. Figure S1** Structure of APY7 and comparison to related APYs

**(A)** Upper panel; schematic drawing of the APY7 protein and genomic DNA, with the protein coding sequence (indicated in grey) interrupted by an intron. qRT-PCR was performed with two primer pairs, located in the 5' and 3' UTRs, respectively. Total RNA was extracted from wild type (Col), *lrx1, lrx1 rol16*, and *lrx1 apy7-1* mutant seedlings. The lower panel shows quantification of RNA levels with the wild type set to 1. **(B)** Comparison of APY7 with APY1,2,6 of Arabidopsis and a human NTPDase. APY6 is the only Arabidopsis APY, besides APY7, to have two TMDs. APYs have five conserved motifs (apyrase conserved regions, ACRs), in APY7 corresponding to positions 150-157 (ACR1), 236-246 (ACR2), 281-293 (ACR3), 312-319 (ACR4), and 566-569 (ACR5). The amino acid sequences of the ACRs are listed. APY7 has a C-terminal extension not found in any other APY protein in Arabidopsis, the biological significance of which remains elusive.

Suppl. Figure S2

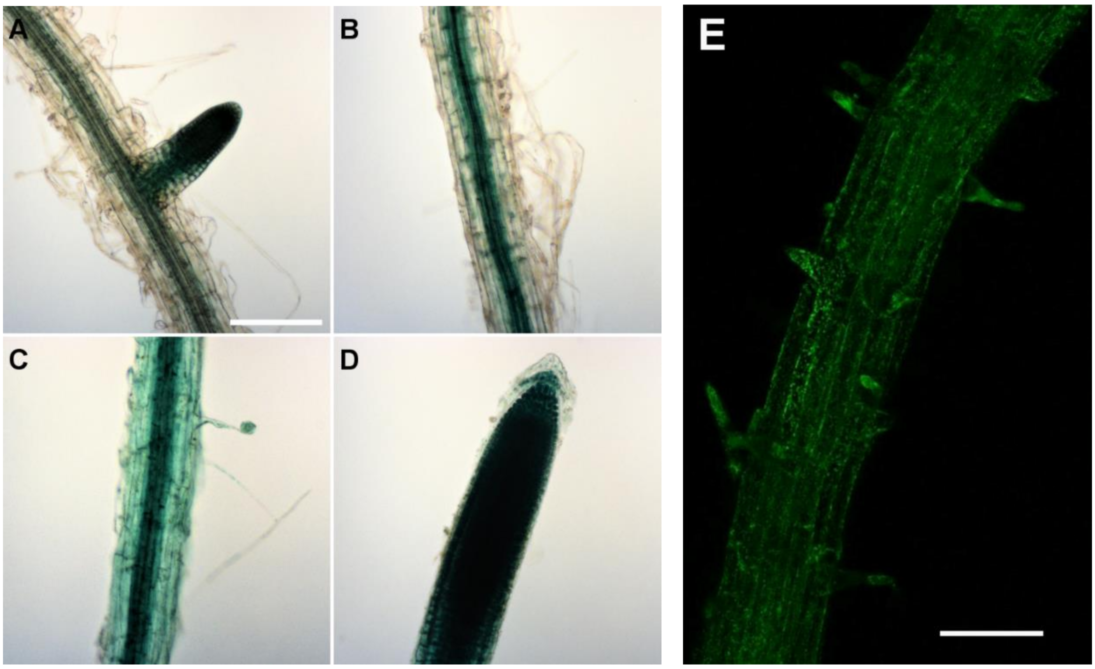


**Suppl. Figure S2** Expression of *APY7* in roots of *Arabidopsis thaliana*.

**(A-D)** *APY7::GUS* fusion construct transformed into Col-0 for expression analysis. The 6-days-old seedlings were stained with GUS for 3-4 hours and the reaction was stopped by adding 70 % ethanol. *APY7::GUS* activity was observed in the root tissue. GUS staining was observed at the lateral root initiation site **(A)**, in the vasculature of the maturation/ differentiation zone **(B)**, diffused expression in the elongation or cell division zone **(C)**, and high expression in the root tip **(D)**. **(E)** GFP fluorescence induced by an *APY7::APY7-GFP* construct reveals expression in all cells with a punctate structure likely representing the Golgi. Bar = 400 µm (A-D have the same magnification).

Suppl. Figure S3

**A**

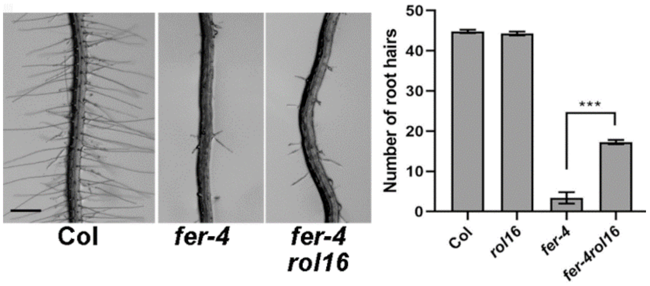


**B**

| parental genotype | total number of progeny analyzed | number of homoz. mutant progeny | frequency of homoz. mutant progeny [%] |
|---|---|---|---|
| *fer-4* +/- | 800 | 60 | 7.5 |
| *rol16* +/- | 840 | 190 | 22 |
| *rol16* +/+; *fer-4* +/- | 800 | 12 | 1.5 |

**C**

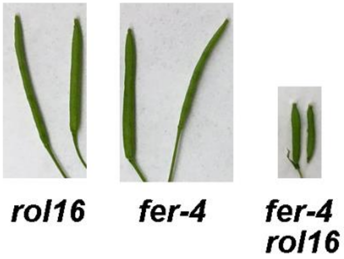


**Suppl. Figure S3** Interaction of the *rol16* and *fer* mutations.

**(A)** *rol16* alleviates the root hair developmental defect induced by the *fer-4* knock out mutant. With more root hairs successfully entering the elongation phase, while remaining shorter than in the wild type. Bar = 500 µm **(B)** The frequency of homozygous mutant *fer-4* is strongly reduced by five-fold in the *rol16* mutant background compared to a wild-type *ROL16* background, suggesting a genetic interaction between *rol16* and *fer-4*. **(C)** Siliques of *fer-4 rol16* double mutants are shorter and contain less seed than the respective single mutants.

Suppl. Figure S4

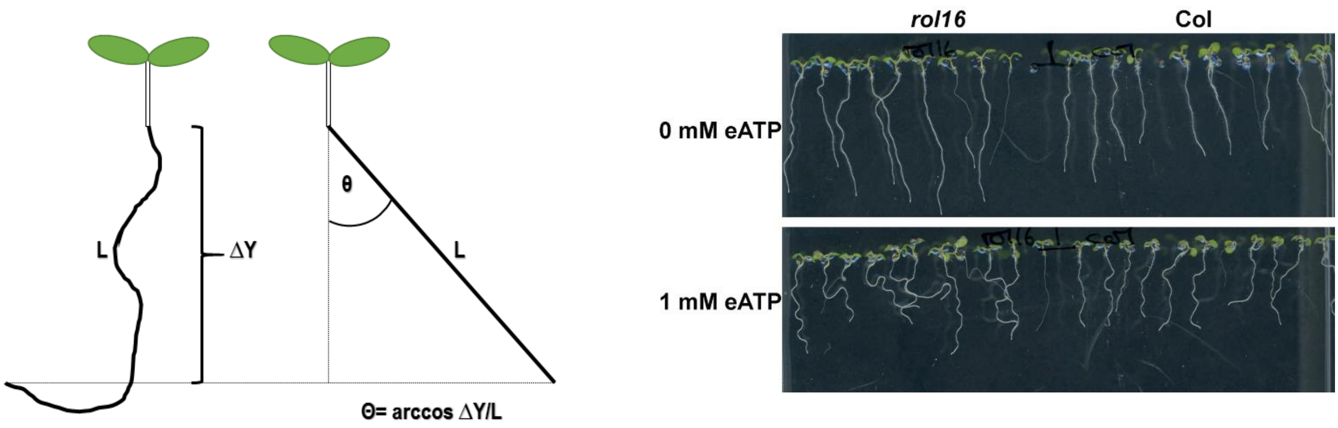


**Suppl. Figure S4** Quantification of vertical growth.

Schematic representation on the left: the total length of the seedlings root (L) and the progression on the Y-axis (ΔY) describe the arccosinus of Θ, a mathematical description of growth behaviour (Grabov et al, 2005). Stronger agravitropic growth or increased root scewing results in an increase in angle Θ. Right: example of Col versus *rol16* mutant seedlings grown with 0 mM ATP (top) and 1 mM ATP (bottom), used for data acquisition. It reveals that with eATP treatment, values are bigger in *rol16* mutants compared ot the wild type.

Suppl. Figure S5

**A**

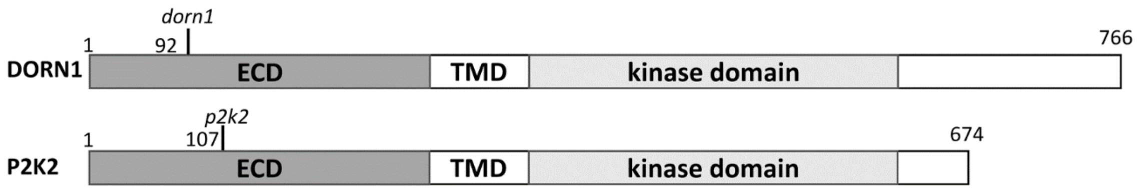


**B**

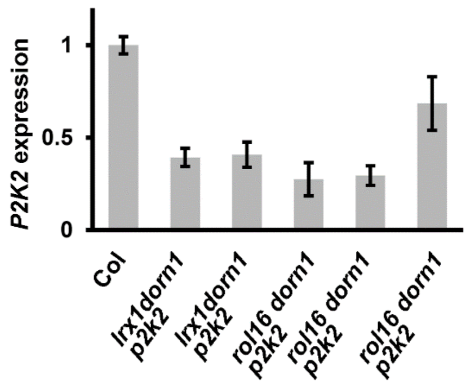


**Suppl. Figure S5** DORN1 and P2K2 protein structure.

**(A)** *P2K1*/*DORN1* (At5g60300) and *P2K2* (At3g45430) encode lectin-like receptor kinases that bind ATP. Numbers indicate amino acid positions, the mutants being used in this work are indicated, the *dorn1* mutant is the T-DNA insertion line SALK_042209 with the insertion site corresponding to amino acid codon 92, *p2k2* represents *CRISPR/CAS9* -induced mutations at the codon 107 that change the reading frame and terminate translation after around 30 amino acids. Numbers indicate amino acid positions. **(B)** *P2K2* expression levels were determined by qRT-PCR on RNA samples from wild-type Col and different *p2k2* alleles, with Col arbitrarily set to 1.

Suppl. Figure 6

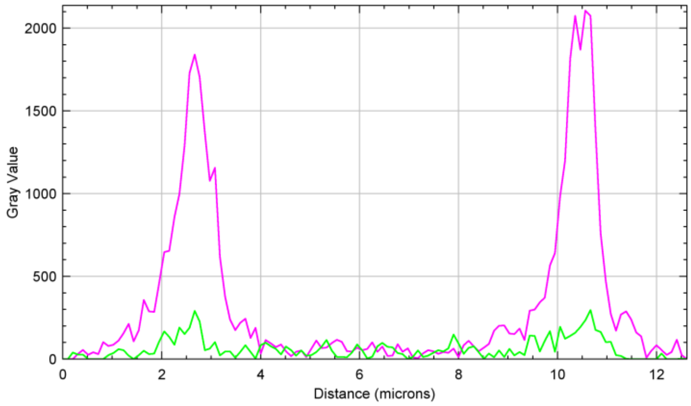


**Suppl. Figure S6** APY7 is not a genuine plasma membrane protein.

Plants expressing APY7-GPF with the plasma membrane marker LTI6b-RFP were used for colocalization. A virtual section through cells revealed strong RFP signal at the cell periphery due to LTI6b-RFP fluorescence in the plasma membrane. By contrast, no signal beyond background could be identified with the GFP filter identifying APY7-GFP.
